# MutagenesisForge: A Framework for Modeling Codon-Level Mutational Biases and Calculating dN/dS

**DOI:** 10.1101/2025.08.28.667654

**Authors:** Cooper Koers, Rob Bierman, Huixin Xu, Joshua M. Akey

## Abstract

The ratio of nonsynonymous (d_N_) to synonymous (d_S_) substitutions in protein-coding genes is a fundamental metric in molecular evolution to test hypotheses about the relative contributions of genetic drift and natural selection in shaping patterns of protein divergence (Williams et al., 2020). However, interpretation of d_N_/d_S_ ratios may be confounded by sequence context and specific substitution models (Hughes, 2007; Kryazhimskiy & Plotkin, 2008). We present MutagenesisForge, a modular command-line tool and Python package for simulating codon-level mutagenesis and calculating d_N_/d_S_ under user-specified conditions. At its core is the MutationModel interface which supports specific substitution matrices and ensures consistency across both Exhaustive and Contextual modes of simulation. These modes allow for users to test evolutionary hypotheses or to generate null distributions of d_N_/d_S_ across a range of biologically relevant models. As large-scale DNA sequencing data sets continue to be generated both within and between species, MutagenesisForge offers a flexible platform for evolutionary analysis and hypothesis testing of mutational processes in protein-coding genes.

## INTRODUCTION

Point mutations that change the DNA sequence in protein-coding regions of the genome are typically classified as synonymous, nonsynonymous (or missense), or protein-truncating (or nonsense) depending on their functional consequences (Nei & Gojobori, 1986). Synonymous mutations do not change the amino acid sequence of a protein whereas nonsynonymous mutations do result in an amino acid change. Protein-truncating mutations introduce a stop codon that results in translation of the protein to prematurely end. Although protein-truncating mutations can have large functional consequences on a protein, here we focus on synonymous and nonsynonymous mutations, which are more abundant in catalogs of natural protein-coding sequence variants and because they have become a cornerstone in evolutionary analyses.

Specifically, a powerful approach for understanding protein evolution is to align a protein-coding DNA sequence of interest from two or more species and calculate the ratio of observed nonsynonymous to synonymous mutations (also referred to as substitutions). This simple, yet fundamentally important measure, provides insight into the selective pressures that a protein has experienced (Williams et al., 2020). More formally, the ratio of nonsynonymous to synonymous mutations is denoted as d_N_/d_S_, where d_N_ is the rate of nonsynonymous mutations normalized by the number of nonsynonymous sites in a protein and d_S_ is the rate of synonymous mutations normalized by the number of synonymous sites in a protein (Nei & Gojobori, 1986). A d_N_/d_S_ = 1 suggests neutral evolution, d_N_/d_S_ < 1 implies purifying selection, and d_N_/d_S_ > 1 is consistent with the effects of positive selection (Kryazhimskiy & Plotkin, 2008). Comparing d_N_/d_S_ ratios across protein-coding genes and species provides considerable insights into the types of proteins and biological processes that have been adaptive in the evolutionary history of distinct species (Bustamante et al., 2005; Nielsen et al., 2005). Despite d_N_/d_S_ ratios providing a powerful metric for detecting selection, a number of factors can complicate its interpretation (Hughes, 2007; Kryazhimskiy & Plotkin, 2008). For example, the substitution model (see below) and sequence context that mutations occur can lead to very different estimates (Goldman & Yang, 1994). While multiple applications exist to perform a d_N_/d_S_ analysis, they do not offer comparative models for individual inputs or the ability to account for evolutionary context. Without context, the value of this ratio may lack definitive biological meaning.

Despite the availability of tools to calculate d_N_/d_S_ values, popular methods lack a modular framework to generate simulated datasets tailored to empirical data (Pond & Frost, 2005, 2005; Yang, 2007; Zhang et al., 2006). To address this limitation, we developed MutagenesisForge, which is a command-line tool that generates simulated sequence data and d_N_/d_S_ values for a wide variety of mutational models. MutagenesisForge allows for the construction of flexible codon-specific null models of the evolution of protein-coding DNA sequences. Thus, MutagenesisForge complements existing software whose functionality is restricted to calculating d_N_/d_S_ values of sequences without the ability to simulate arbitrarily complex context-dependent mutations. With MutagenesisForge, simulation functionality is brought to the user directly through either a command line interface (CLI) or as an installable python package for programmatic use. We anticipate MutagenesisForge will have many uses in comparative genomic and evolutionary analyses.

## METHODS

### Evolutionary Model-Dependant Substitution for Codon-level Mutational Biases

MutagenesisForge provides a platform to simulate nucleotide-level substitutions according to user-selected evolutionary models. The models are shared with the exhaustive and context-specific methods, allowing for comparisons. Given an input nucleotide sequence of a protein of interest and a substitution model with given parameters, a corresponding nucleotide is returned following the probabilities of the substitution matrix. A list of substitution models implemented in MutagenesisForge is summarized in Figure 1.

**Figure 1:**
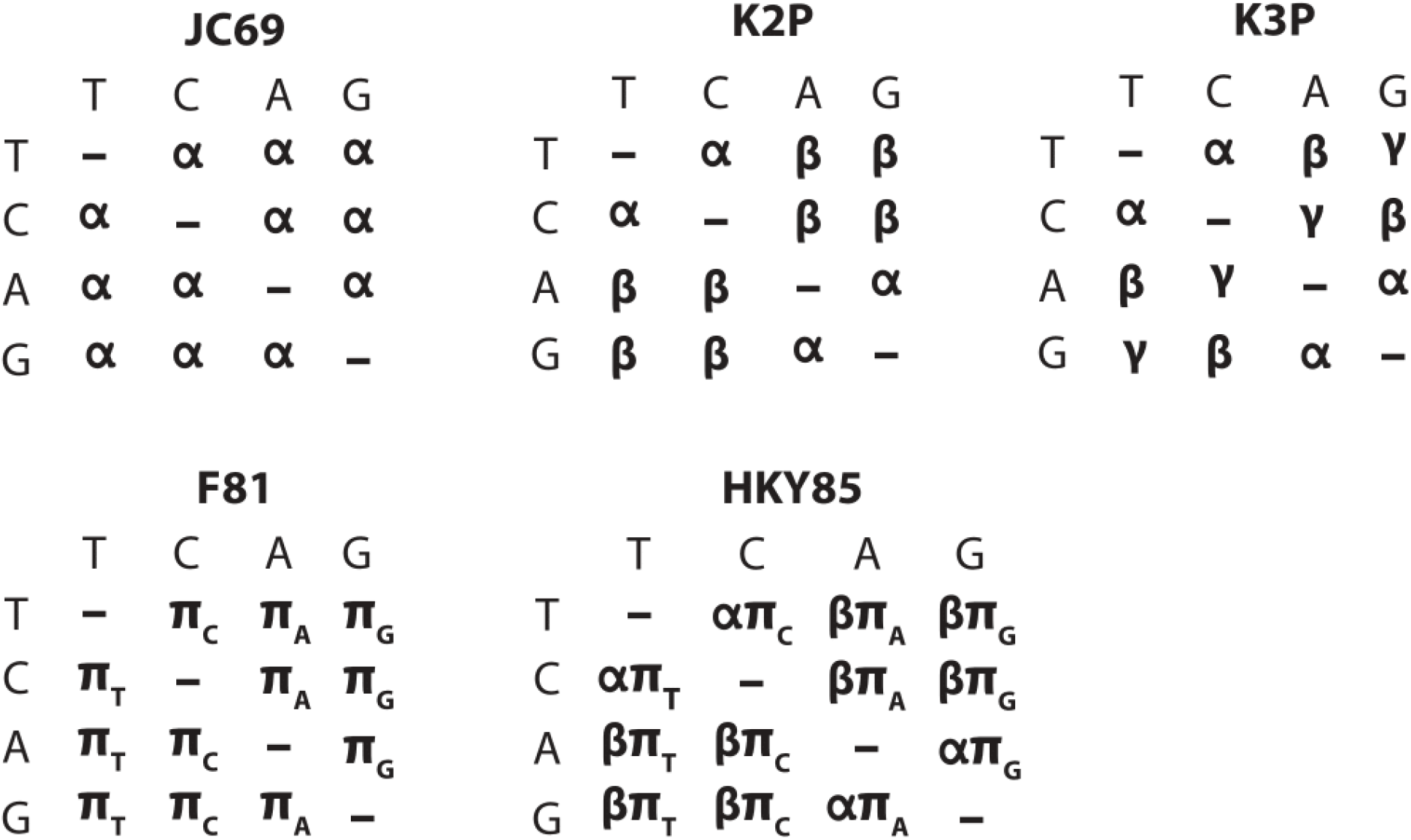
Matrix representations of supported evolutionary substitution models. MutagenesisForge supports the Kimura 2-parameter (K2P), Kimura 3-Parameter (K3P), Jukes-Cantor (JC69), Felenstein 4-parameter (F81), and Hasegawa, Kishino, and Yano (HKY85) substitution models of evolution to simulate single-nucleotide substitutions (Felsenstein, 1981; Hasegawa et al., 1985; Jukes & Cantor, 1969; Kimura, 1980, 1981). The rows of each matrix represent the existing nucleotide and columns denote the mutated nucleotide. The nucleotides are abbreviated as T (thymine), C (cytosine), A (adenine), and G (guanine). Thus, the element in a substitution matrix corresponding to a row with C and column with G indicates a mutation from C to G. Symbols within the matrix denote mutation rates. For example, in the JK69 model, the mutation rate from one nucleotide to another is □ for all possible substitutions whereas in the K2P model the mutation rates of transitions (C↔T and A↔G) and transversion (C↔A, C↔G, A↔T, and G↔T) are different and occur at rates denoted as □ and □ respectively.

### Approaches for Simulating d_N_/d_S_

MutagenesisForge offers two distinct approaches to simulate data that we refer to as **Exhaustive** and **Contextual**. Both of these modes allow comparison between empirical values of d_N_/d_S_ and simulated data derived from user-specified parameters and a desired evolutionary model. Below, we describe each simulation mode in more detail.

#### Exhaustive Mode

The exhaustive mode of MutagenesisForge systematically simulates all possible single nucleotide substitutions across a protein-coding DNA sequence. The input format for the protein coding sequence of interest is a standard FASTA file. Starting from the first identified start codon, the algorithm consists of a sliding window analysis incrementing by codon, where each position has all possible mutations made and categorized as either synonymous or nonsynonymous. These data are stored and used as inputs to calculate d_N_/d_S_. This mode accepts FASTA files of the protein-coding sequences that the user is interested in and supports user-defined evolutionary substitution models. Under specific substitution models, each exhaustive mutation is weighed according to its probability of occurrence. Users can restrict the exhaustive search to specific genomic regions of interest by setting an optional flag. The workflow of determining the number of missense and synonymous sites per codon is presented in Figure 2.

**Figure 2:**
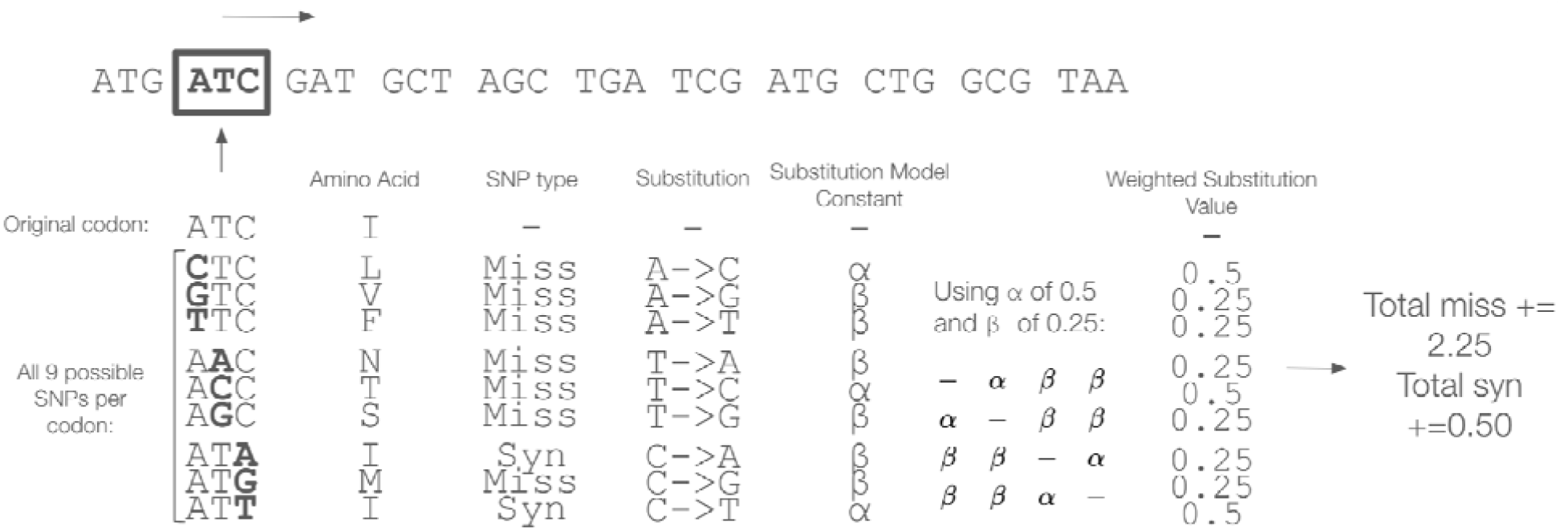
Overview of Generating d_N_/d_S_ Values using the Exhaustive Mode of MutagenesisForge with the K2P Substitution Model. The schematic illustrates how MutagenesisForge calculates the number of missense and synonymous sites per codon in accordance with the respective substitution model.

#### Context Mode

It is well known that mutation rates can be context-specific (Aggarwala & Voight, 2016). Specifically, the rate of a particular mutation can differ depending on the specific nucleotides surrounding the focal nucleotide site. For instance, the rate of a C>T mutation can vary dramatically depending on the specific nucleotides immediately up and downstream of it (i.e., the rate of C**C**G > C**T**G mutations can be markedly different compared to the rate of A**C**T > A**T**T mutations even though both mutations involve a C changing to a T). The context mode of MutagenesisForge non-deterministically simulates mutations based on the nucleotide context surrounding variants in a VCF file by forming codons from bases flanking the nucleotides of interest from the reference FASTA file, where the position of the initial variant is randomized in the newly-formed codon (Aggarwala & Voight, 2016). The initial variant is then mutated according to the mutation model provided. The workflow of the context mode is portrayed in Figure 3, highlighting how the process acts on each variant. Analysis between the mapped codon and mutated mapped codon then allows for the calculation of a null model of d_N_/d_S_ from the lens of the nucleotide context of the input file.

**Figure 3:**
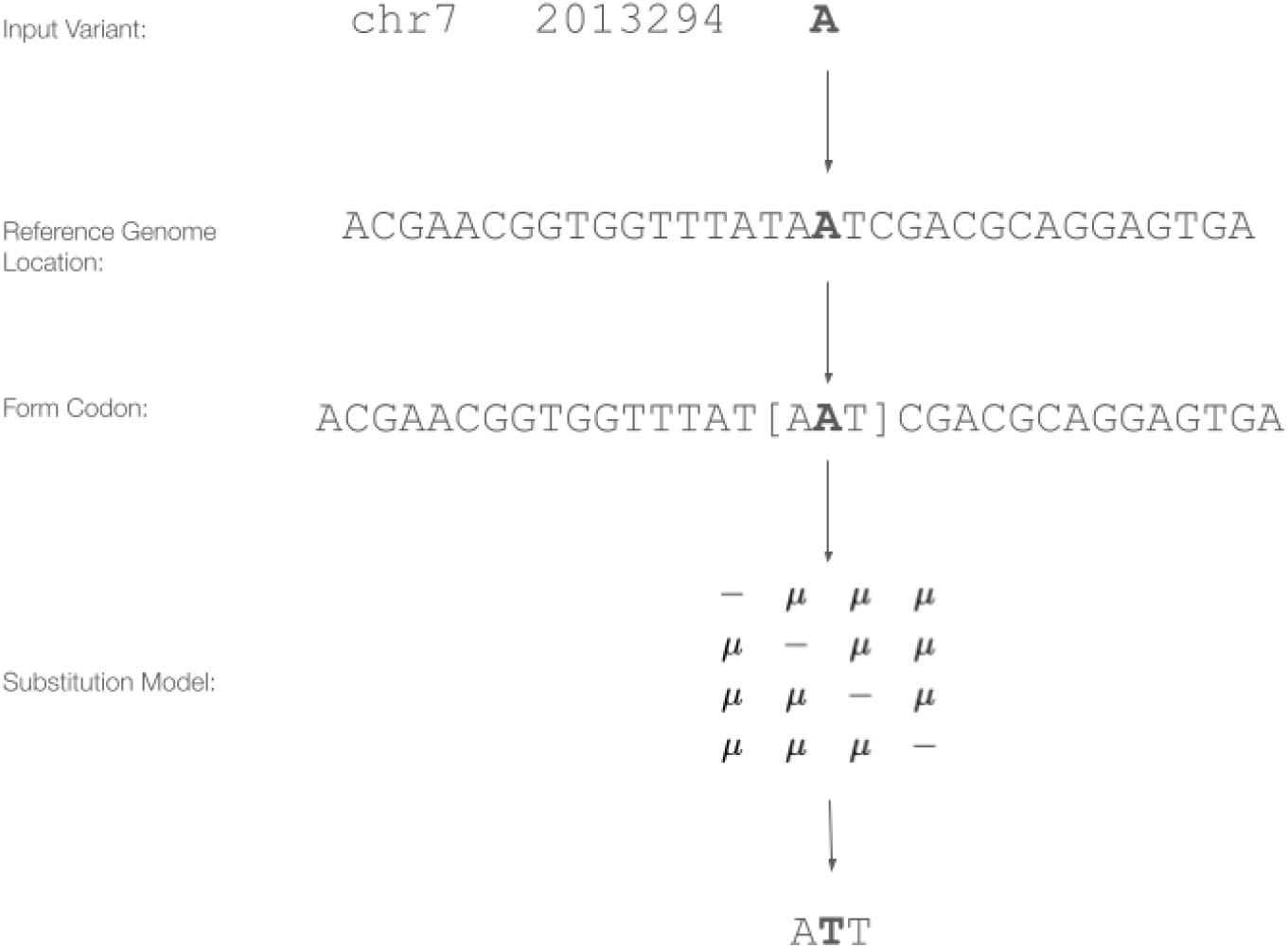
Calculating d_N_/d_S_ values using the Context Mode of MutagenesisForge with the JC69 Substitution Model. Given the input variant, a match is found within the coordinates of interest. From this coordinate, a codon is constructed from flanking bases, with the variant of interest at a random position within the codon. A substitution is made according to the input substitution model. Data from each substitution is then used to calculate the d_N_/d_S_ value of the simulation to return a context-aware simulated value.

## APPLICATIONS AND RESULTS

### Exploratory Results

MutagenesisForge was used to generate null model d_N_/d_S_ values for the *BRCA1* human gene. *BRCA1* encodes the BRCA1 protein, which is crucial for DNA repair and genome stability pathways, causing *BRCA1* to be among the most well-studied tumor suppressor genes in the annotated human genome (Zhong et al., 2023). To simulate a null model using the human reference sequence for *BRCA1*, we used MutagenesisForge in both context and exhaustive mode with different substitution models. Each substitutional model supported by MutagenesisForge was used, including models that support differential transversion and transition ratios (K2P, K3P, and HKY85). For each of these models, two matrices were tested: one that biased transitions and one that biased transversions. Using MutagenesisForge exhaustive mode, each substitution model was tested on the *BRCA1* coding sequence. Results of these deterministic analyses are presented in Table 1.

**Table 1:** Simulating d_N_/d_S_ values using MutagenesisForge Exhaust for Null Model Analysis of *BRCA1* in Humans. *BRCA1* d_N_/d_S_ values were benchmarked using MutagenesisForge exhaustive mode.

| Substitution Model | $d_N/d_S$ |
| --- | --- |
| K2P transition bias | 0.825 |
| K2P transversion bias | 1.481 |
| K3P transition bias | 0.958 |
| K3P transversion bias | 1.623 |
| HKY85 transition bias | 0.852 |
| HKY85 transversion bias | 1.489 |
| JC69 | 1.000 |
| F81 | 1.026 |

Seen in the results are increased d_N_/d_S_ values specifically for substitution models with transversion biases, a confirmatory result. On average, transversion substitutions result in nonsynonymous mutations at higher rates than transition substitutions. Thus, data enriched for transversion substitutions are expected to exhibit higher d_N_/d_S_ values.

For each parameter combination, we performed 200 simulations using MutagenesisForge context to create distributions of d_N_/d_S_ values. The simulated null model distributions from context mode are seen in Figure 4.

**Figure 4:**
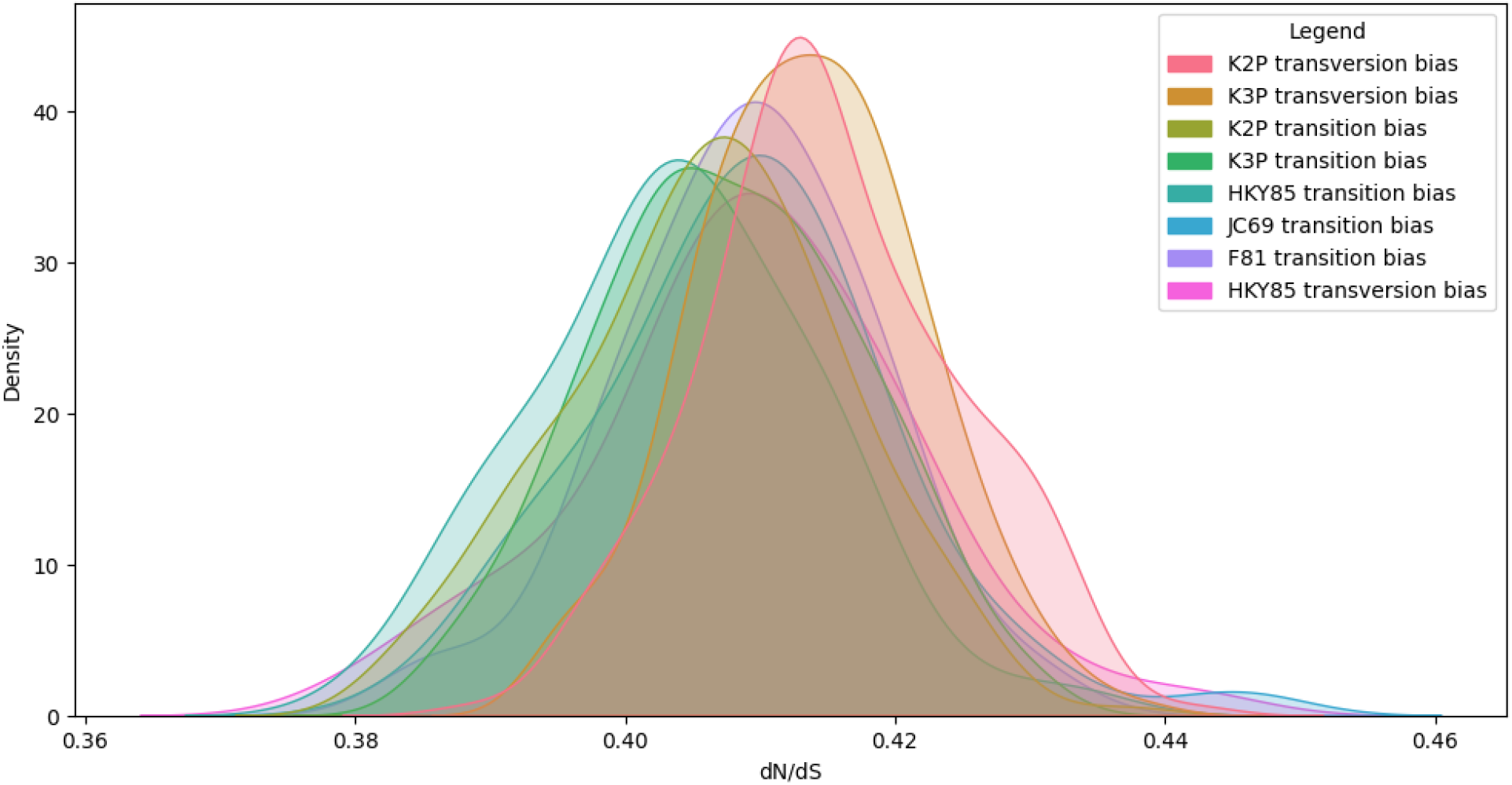
Simulating d_N_/d_S_ values using MutagenesisForge Context Mode for Null Model Analysis of *BRCA1* in Humans Yields Consistently Low Values. *BRCA1* d_N_/d_S_ values were benchmarked using MutagenesisForge exhaustive mode. Distributions returned are plotted by the substitution model used.

Analysis of *BRCA1* context mode simulated d_N_/d_S_ distributions reveals an insignificant dependency (p-value > 0.05) on the substitution model selected. However, comparison between exhaustive and context mode outputs demonstrates that the sequence composition of *BRCA1* leads to a higher synonymous substitution rate relative for random mutations compared to nonsynonymous substitutions (Figure 4). These results suggest that sequence composition can influence the null distribution of d_N_/d_S_.

### Applications in Research

MutagenesisForge was used in Xu et al. 2025 to investigate mutational models that were consistent with empirical patterns of d_N_/d_S_ caused by protein-coding somatic mutations. Specifically, high-coverage exome sequencing was performed on 265 tissues from 14 individuals (average of 19 tissues per individual) and this data was used to identify somatic mutations. The compendium of somatic mutations resulted in a large excess of nonsynonymous compared to synonymous change. Thus, we used MutagenesisForge to generate mutations for a variety of mutational models and compared the mutational spectrum and distribution of d_N_/d_S_ between the empirical and simulated datasets (Xu et al. 2025). MutagenesisForge enabled us to make statistically rigorous inferences of a complicated biological phenomenon that would not have been possible without a flexible tool to simulate mutations in protein-coding DNA sequences under a wide variety of mutational models.

As DNA sequencing datasets continue to be generated at a frenetic pace, we anticipate MutagenesisForge will be a valuable tool to interpret the evolutionary forces shaping patterns of protein-coding sequence variation within and between populations and species.

## ACKNOWLEDGEMENTS

We would like to thank members of the Akey laboratory for helpful feedback related to this work and for comments on the manuscript. This work was supported in part by NIH grants U01HG007591 and R01GM110068 to JMA.

